# Processing and evaluating the quality of genome-wide nascent transcription profiling libraries

**DOI:** 10.1101/2022.12.14.520463

**Authors:** Thomas G. Scott, André L. Martins, Michael J. Guertin

## Abstract

Precision genomic run-on assays (PRO-seq) quantify nascent RNA at single nucleotide resolution with strand specificity. Here we deconstruct a recently published genomic nascent RNA processing pipeline (PEPPRO) into its components and link the analyses to the underlying molecular biology. PRO-seq experiments are evolving and variations can be found throughout the literature. The analyses are presented as individual code chunks with comprehensive details so that users can modify the framework to accommodate different protocols. We present the framework to quantify the following quality control metrics: library complexity, nascent RNA purity, nuclear run-on efficiency, alignment rate, sequencing depth, and RNA degradation.

## 1 Introduction

Genomic nascent RNA profiling assays, such as precision genomic run-on (PRO-seq) ***(1)***, quantify the precise position and direction of transcriptionally engaged RNA polymerases. Quantifying nascent RNA complements conventional RNA-seq by directly measuring active transcription. Steady-state RNA levels are influenced by RNA stability, so we can leverage the discordance between RNA-seq and PRO-seq expression to estimate genome-wide RNA half-lives ***(2)***. Bidirectional transcripts are a hallmark of enhancers and promoters. We can detect these short and unstable transcripts with PRO-seq to directly infer regulatory element activity as an orthogonal approach to chromatin accessibility assays ***(3, 4)***. Similarly to regulatory elements, gene isoforms can vary between cell types and conditions. We can use RNA-seq to define splice variants and PRO-seq to identify differing primary transcript boundaries ***(5)*** and transcription start sites ***(6)***. Additionally, PRO-seq sensitively detects immediate changes in transcription without the need for mature RNAs to accumulate or degrade. Lastly, nascent RNA profiling determines RNA polymerase density within all genomic features, such as promoter-proximal regions, gene bodies, and enhancers ***(7–9)***. Changes in RNA polymerase distribution within these regions can inform on how various treatments and stimuli regulate steps in the transcription cycle ***(9–11)***. Here, we describe quality control metrics that are used to determine if PRO-seq libraries are worth proceeding with these or other downstream analyses. New genomic nascent RNA-seq methodologies ***(4, 12–15)*** necessitate flexible analysis workflows and standardized quality control metrics ***(16)***. We present the workflow as deconstructed code that can be adapted to fit a diversity of protocols and experimental details.

## 2 Software and hardware requirements

Many processes, downloads, and software installations are reused throughout the analyses. Users should periodically check for updated annotations and new software releases.

### 2.1 Dependencies, software, and scripts

We present specialized software and scripts herein, but much of the workflow depends upon more general software. These general bioinformatic software tools are well-maintained and documented, so we provide short descriptions and the links below.

`bedtools`: a comprehensive suite of tools that efficiently perform a wide range of operations on genomic intervals. https://bedtools.readthedocs.io/en/latest/ ***(17)***
`bowtie2`: aligns sequencing reads to reference sequences. http://bowtie-bio.sourceforge.net/bowtie2/index.shtml ***(18)***
`cutadapt`: removes a defined sequence, such as adapter sequence, from sequencing reads. https://cutadapt.readthedocs.io/en/stable/ ***(19)***
`fastq_pair`: outputs only sequencing reads that have a matched paired end read. https://github.com/linsalrob/fastq-pair ***(20)***
`FLASH`: merges paired end reads by detecting overlapping sequence. https://ccb.jhu.edu/software/FLASH/ ***(21)***
`fqdedup`: removes duplicated sequences from FASTQ files. https://github.com/guertinlab/fqdedup ***(22)***
`samtools`: a suite of tools for parsing and interfacing with high throughput sequencing data files. http://www.htslib.org ***(23)***
`seqOutBias`: software that parses files and outputs desired formats with the option to correct enzymatic sequence biases. https://github.com/guertinlab/seqOutBias ***(24)***
`seqtk`: a multifunctional toolkit for processing sequence files, including trimming a defined number of bases from the ends of reads and reverse complementing sequencing reads. https://github.com/lh3/seqtk ***(25)***
`sratoolkit`: a suite of tools that interface with data deposited into the Sequence Read Archive. https://github.com/ncbi/sra-tools
`wget`: retrieves files from a wide range of internet protocols. https://www.gnu.org/software/wget/
`R` packages:
`lattice`: graphics plotting package. https://cran.r-project.org/web/packages/lattice/lattice.pdf ***(26)***
`DESeq2`: statistical package for quantifying differences in counts-based genomics data. https://bioconductor.org/packages/release/bioc/html/DESeq2.html ***(27)***

In addition, we developed the following software and `R` scripts to facilitate data analysis and graphical output. Below, we use `wget` to retrieve the software and scripts. The command `chmod +x` changes the permissions of the files to executable.

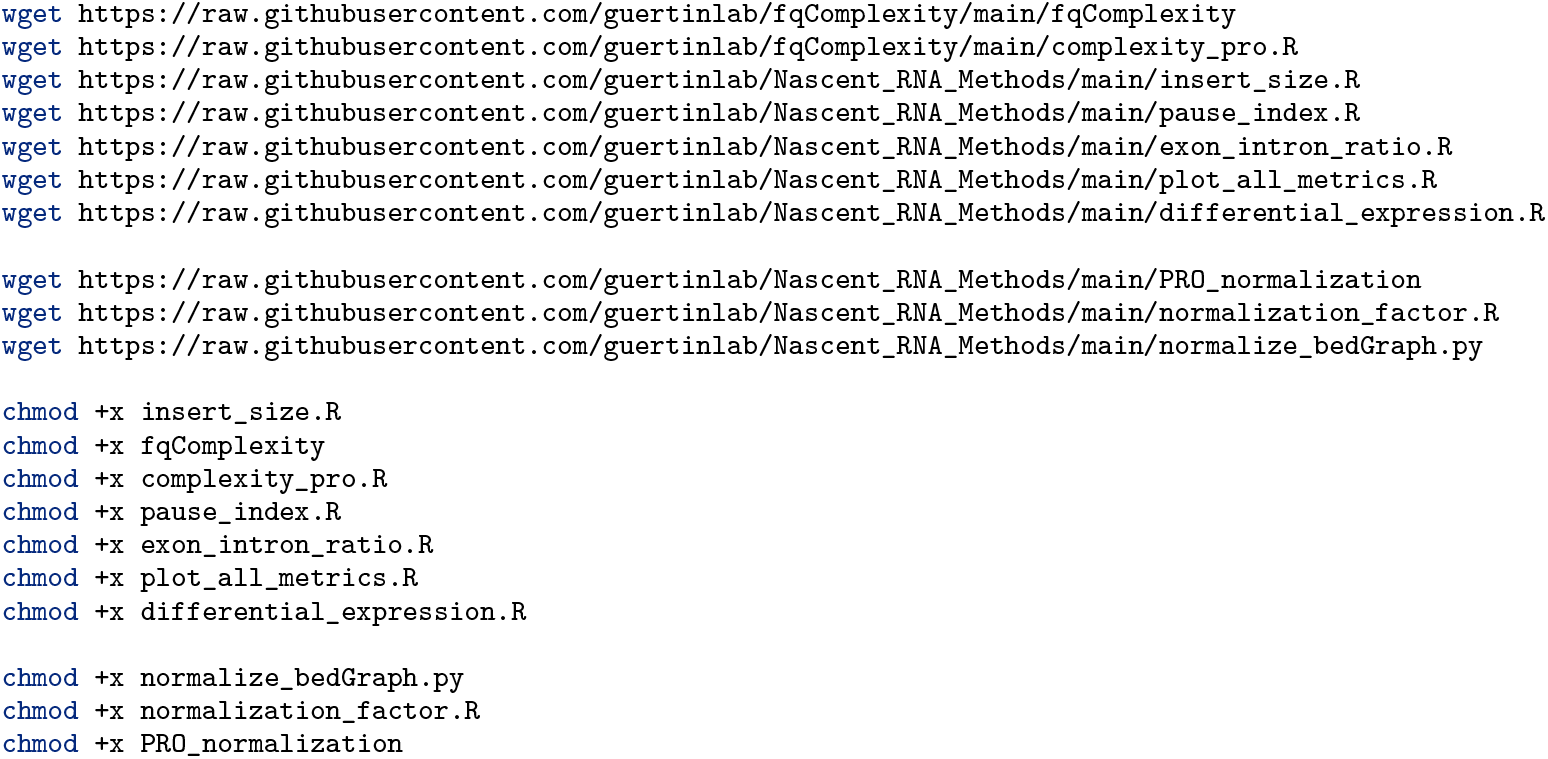

Next, move the software dependencies and `R` scripts to a directory within the `$PATH` variable.

### 2.2 Hardware

This workflow requires a single core computer, 8Gb of RAM, and 200Gb hardrive space. However, more RAM and multiple cores will greatly reduce compute time.

## 3 Genome and annotation downloads and processing

### 3.1 Reference genomes

PRO-seq experiments have been performed in a variety of organisms including yeast ***(28)***, Drosophila ***(1, 8)***, and humans ***(29)***. Analysis of the data requires alignment to a reference genome annotation. The first step is to use `wget` to retrieve the reference genome. Many websites host the assembly data in FASTA format, such as the human genome build 38 shown below retrieved from the UCSC genome browser server ***(30)***. The `gunzip` command unzips the reference genome file and `bowtie2-build` indexes the file to allow for efficient alignment. The code also retrieves, unzips, and builds the human rDNA reference genome ***(31)*** so that we can calculate rDNA alignment rates as a metric for nascent RNA purity.

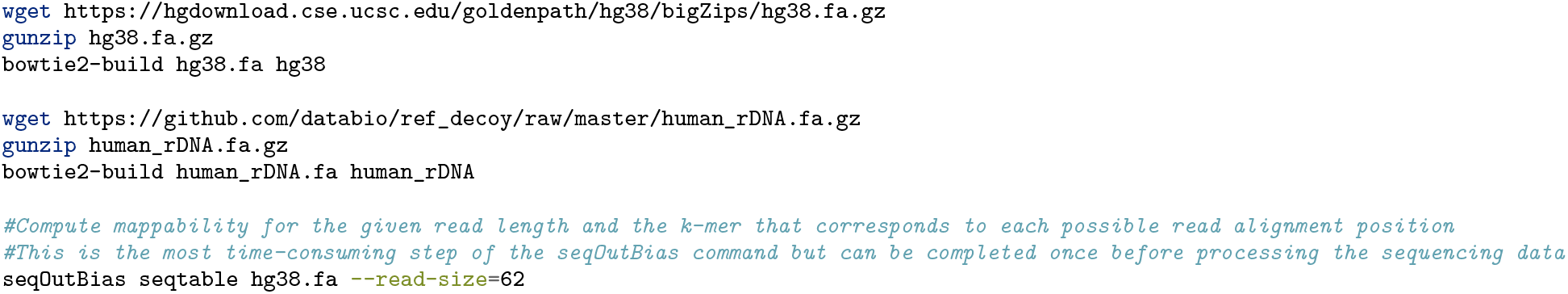

### 3.2 Reference gene annotation

The quality control metrics outlined herein require the counting of sequence reads that align to three genomic features: exons, intron, and promoter-proximal pause regions. Gene annotations are available from many sources and we outline retrieval and parsing of GTF files from Ensembl ***(32)***. The Ensembl website (http://www.ensembl.org/index.html) contains the information for the latest release, which at the time of writing this manuscript is release 104 for hg38. After retrieving and unzipping the file we parse out all exon 1 annotations. These coordinates include all annotated transcription start sites that correspond to different gene isoforms. Ensembl chromosome numbers do not include the preceding “chr”, so the first `sed` command appends “chr” to the chromosome name for downstream compatibility with the reference genome chromosome names. The output is then piped to `awk`, which prints the following fields: chromosome coordinates in columns 1-3, Ensembl transcript ID (ENST), gene name, and strand. Subsequent `sed` commands drop the semicolon and quote characters from the gene and Ensembl IDs while editing the mitochondrial chromosome to match the reference genome, “chrM” replacing “chrMT”. Finally we sort the exon output by the first column (chromosome), then the second column (starting position), in ascending order. The gene annotations are processed similarly, except the Ensembl gene ID (ENSG) replaces ENST in column 4 and we sort the file by the fifth column (gene name).

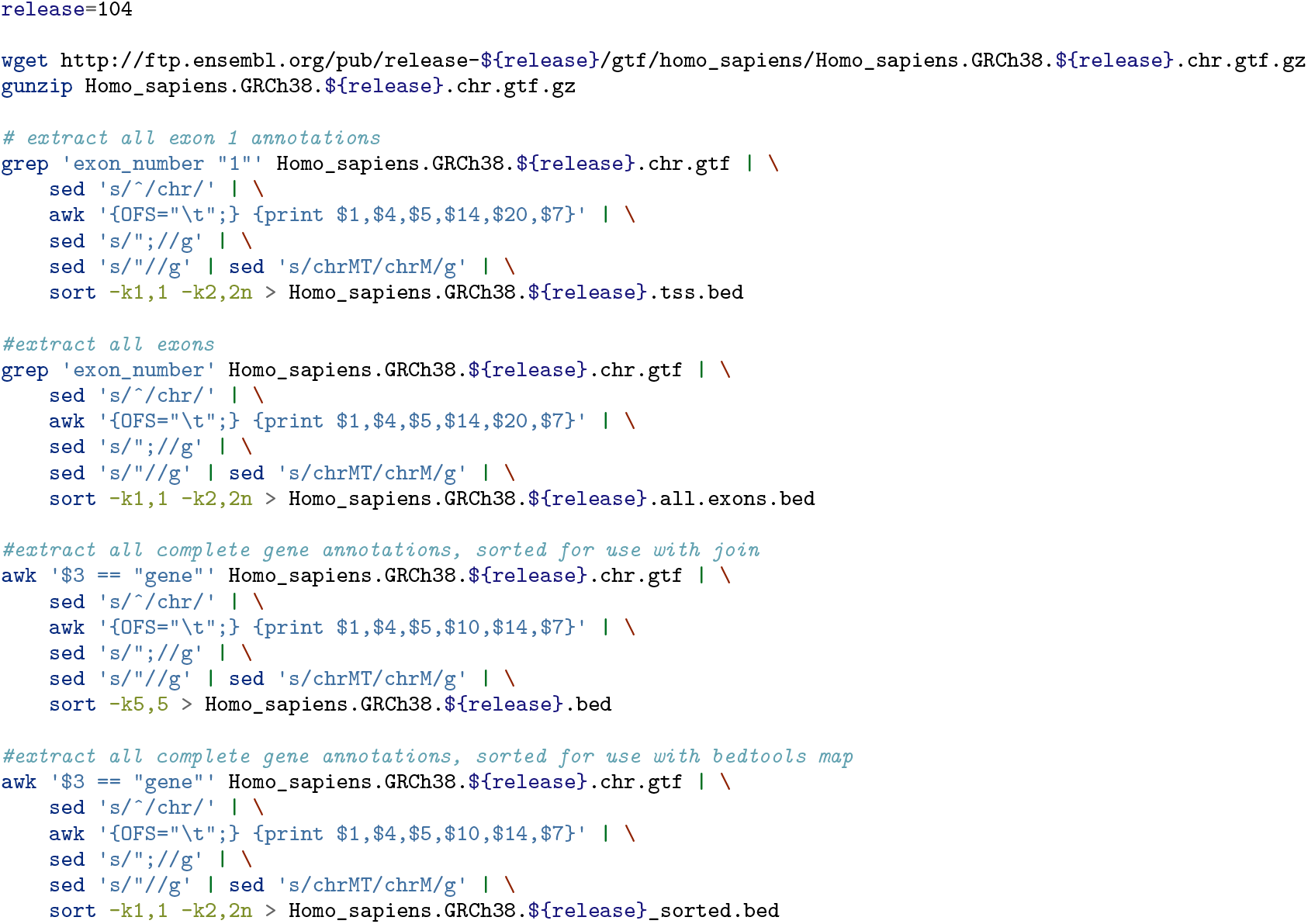

The following operations output: 1) a set of exons that excludes all instances of first exons, 2) all potential pause regions for each gene, and 3) all introns. There are many exon 1 gene annotations depending on gene isoforms and the upstream most annotated TSS is not necessarily the most prominently transcribed isoform. We define the pause window for a gene as position 20 - 120 downstream of the most prominent TSS. The most prominent TSS is determined by calculating the density in this 20 - 120 window for all annotated TSSs for each gene and choosing the TSS upstream of the most RNA-polymerase dense region for each gene.

In order to define these windows, we use `mergeBed` to collapse all overlapping exon intervals and then `subtractBed` to exclude all first exon coordinates from the merged exon file. Since `mergeBed` drops the gene name, we use `intersectBed` to reassign gene names to all remaining exons. The `awk` command defines the 100 base pause region window downstream of all transcription start sites based on the gene strand. Lastly, we subtract the exons from the full gene coordinates to produce the intron annotations.

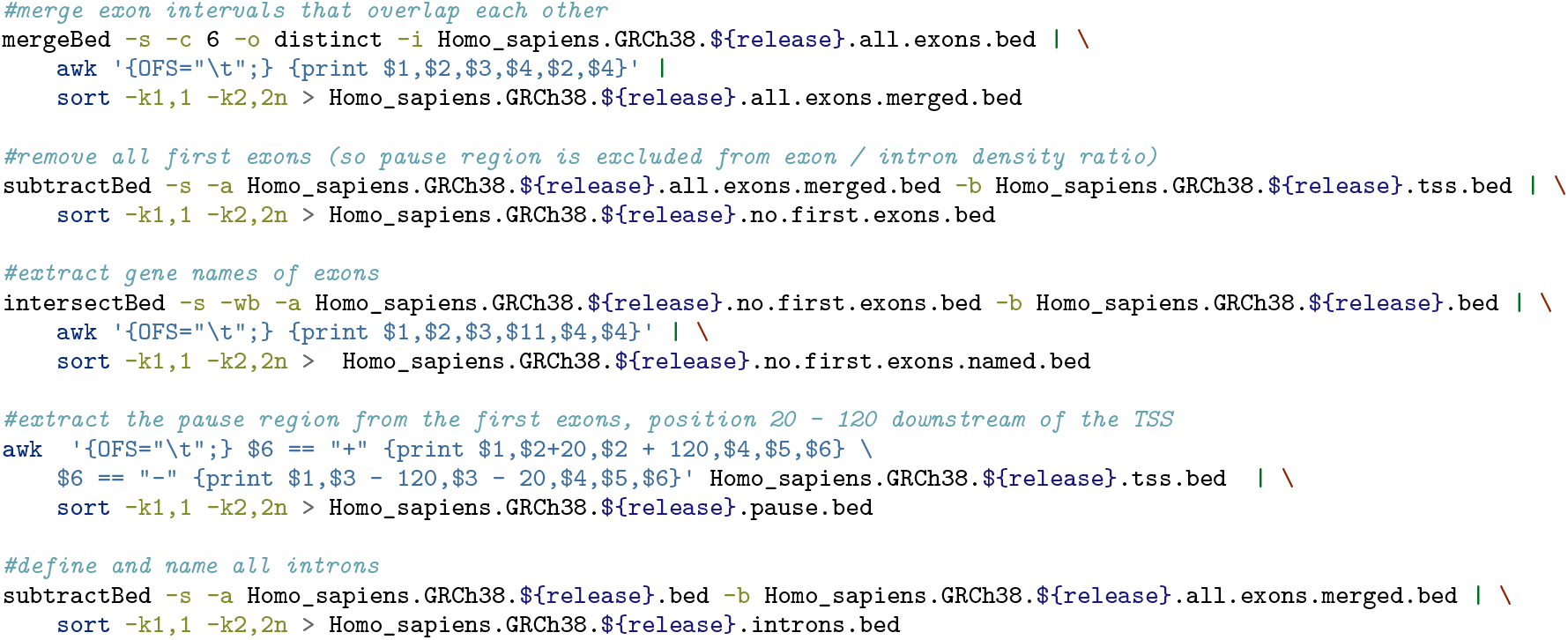

## 4 Processing PRO-seq data

### 4.1 Initialize variables

In order to automate the processing and naming of output files, we conform to a strict naming convention for the FASTQ files: cell-Type_conditions_replicate_pairedend.fastq.gz. For example, a gzipped paired end 1 (PE1) file from the first replicate of treating T47D cells with DMSO would be: `T47D_DMSO_rep1_PE1.fastq.gz`.

We first initialize six variables:

`$directory`: location of the sequencing files
`$filename`: name of the gzipped paired end 1 FASTQ file.
`$annotation_prefix`: Ensembl gene annotation GTF prefix; this is the user-defined prefix from above.
`$UMI_length`: length of the UMI on the 5’ end of the paired end 1 read.
`$read_size`: read length minus UMI length.
`$cores`: number of cores for parallel processing.
`$genome`: absolute or relative path to the genome FASTA file.
`$genome_index`: the basename (including the path) of the genome index files from bowtie2-build.
`$prealign_rdna_index`: the basename (including the path) of the prealign rDNA index files from bowtie2-build.
`$tallymer and $table`: outputs of the seqOutBias command above for a given genome and read length.

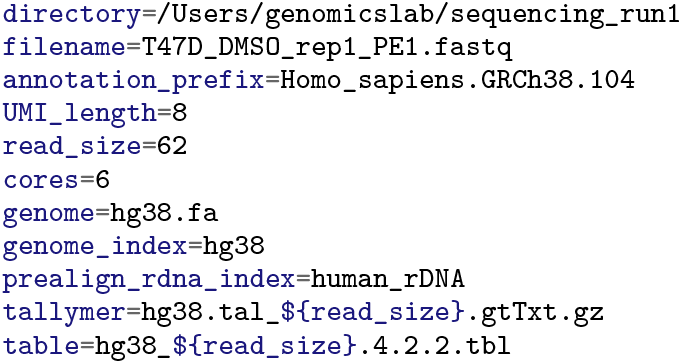

### 4.2 Preprocessing

Make a working directory, download the files from GEO accession GSE184378, and save the basename as a variable.

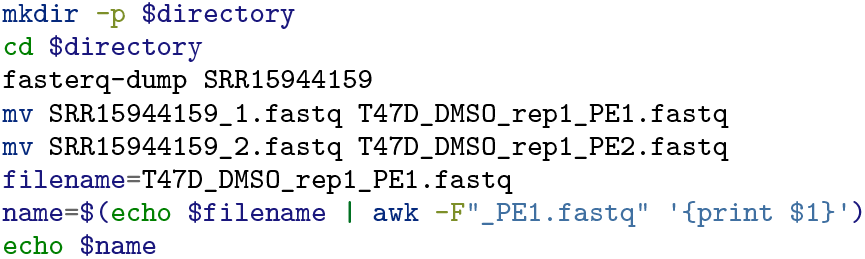

### 4.3 Processing reads

Here we describe processing and analysis of paired end PRO-seq libraries with unique molecular identifiers ligated to the 3’ end of the nascent RNA. The user may need to modify or omit specific steps in order to accomodate their library preparation protocol.

The first processing step is to remove adapter sequences and simultaneously discard reads that have insert sizes of one base. If the adapters ligate directly to one another, the UMI is treated as an insert to `cutadapt` and the effective insert length is equal to the UMI length. The option `-m $((UMI_length+2))` provides a one base buffer and discards reads with a length of less than the UMI length + 2.

The fraction of reads that represent adapter/adapter ligation products is a useful metric to help determine the raw read depth needed to achieve a specified aligned read depth. FASTQ files contain four lines per sequence entry, so we calculate the raw sequencing depth by first using `wc −l` to count the number of lines in the original FASTQ file and using `awk ‘{print $1/4}’` to divide by 4. We perform the same operation on the file containing reads with 0 or 1 base insertions. Finally, we use `$(echo “scale=2; $PE1_w_Adapter / $PE1_total” | bc)` to divide the adapter/adapter ligation product value by the total and round to the nearest hundredth.

The proportion of reads that are adapter/adapter ligation products varies widely depending upon whether a size selection was performed in the library preparation. We recently excluded the size selection step from the PRO-seq protocol to reduce bias against small RNA inserts ***(9, 16)***. If no size selection is performed, the adapter/adapter ligation fraction can be quite high resulting in low effective sequencing depth. In a later section we provide a formula for determining the required raw sequencing depth to result in a desired number of concordant aligned reads. We typically recommend further sequencing if all other QC metrics indicate that the data is high quality. However, if more than 80% of the reads are adapter/adapter ligation products, the user should balance the cost of performing another experiment with sequencing uninformative adapter sequences.

The fraction of adapter/adapter reads and this 0.80 threshold is printed to `${name}_QC_metrics.txt`. We continue to append all metrics and thresholds to this file and plot the data at the end of the workflow.

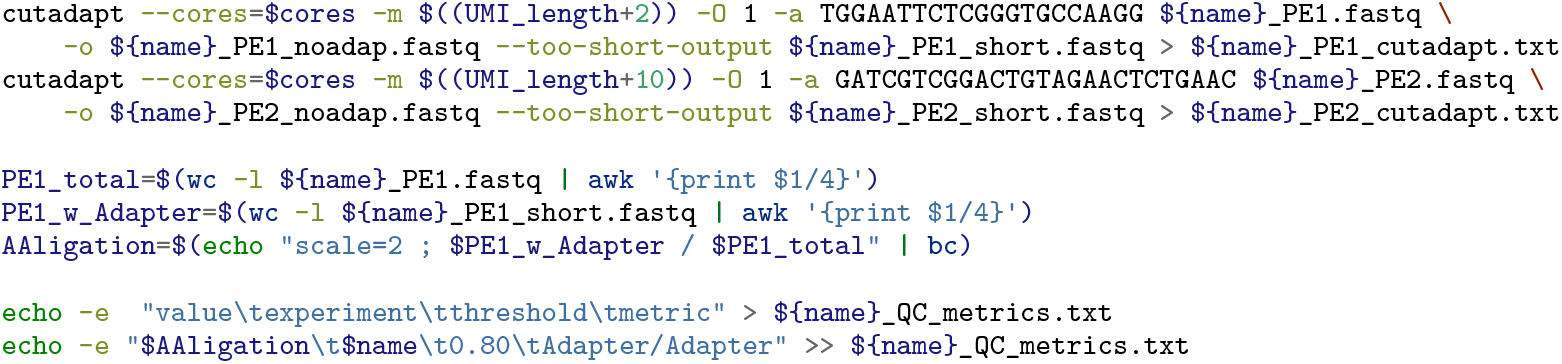

The next step removes reads that are shorter than 10 bases.

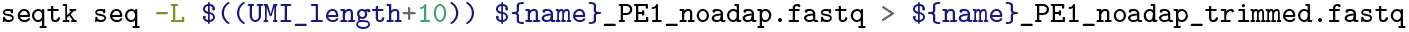

A proportion of short nascent RNAs from different cells are identical because their 5’ end corresponds to a transcription start site, and their 3’ end is located within a focused promoter-proximal pause region ***(1)***. Therefore, we cannot filter potential PCR duplicates based on whether two independent pairs of reads have identical paired end read alignments. We rely on the presence of the UMI to remove PCR duplicates from the PE1 FASTQ file. We use `fastq_pair` to deduplicate the PE2 read by pairing with the deduplicated PE1 file.

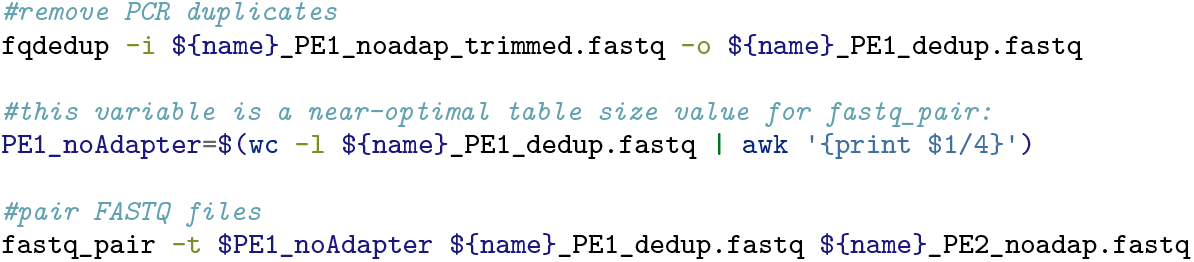

### 4.4 RNA degradation ratio score

An abundance of short inserts within a library indicates that RNA degradation occurred. We measure RNA degradation by searching for overlap between paired end reads with flash and plotting the resultant histogram output with `insert_size.R` (Figure 1). RNA starts to protrude from the RNA Polymerase II exit channel at approximately 20 bases in length, so 20 bases of the nascent RNA is protected from degradation during the run-on. Libraries with a substantial amount of degradation after the run-on step are enriched for species in the range 10 - 20. We empirically found that there are fewer reads within the range of 10 - 20 than within the range of 30 - 40 for high quality libraries ***(16)***. A degradation ratio of less than 1 indicates a high quality library. This metric becomes unreliable if the protocol includes size selection to remove adapter/adapter ligation products. Size selection inevitably removes some small RNAs and inflates this ratio.

**Figure 1:**
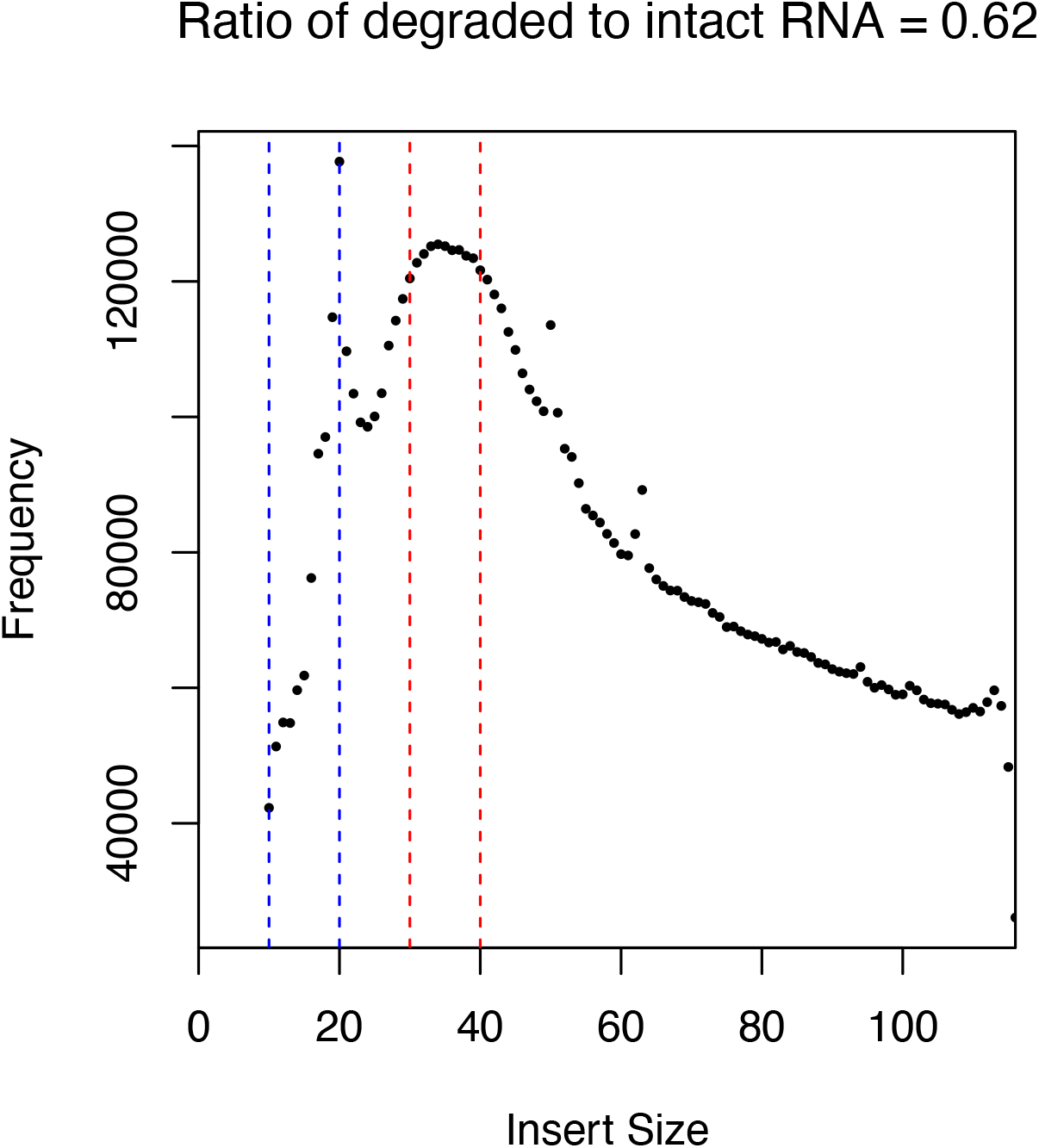
Library insert size is a measure of RNA degradation. The plot illustrates the frequency (y-axis) of insert size lengths (x-axis) for the PRO-seq library. The ratio of read counts in the 10 - 20 base range (blue region) to read counts in the 30 - 40 range (red region) is the degradation ratio. High quailty PRO-seq libraries have degradation ratios less than 1.

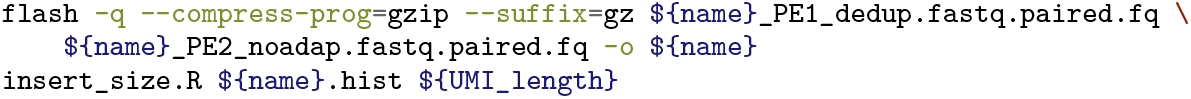

### 4.5 Processing for alignment

The final processing step removes the UMI from and reverse complements both paired end reads.

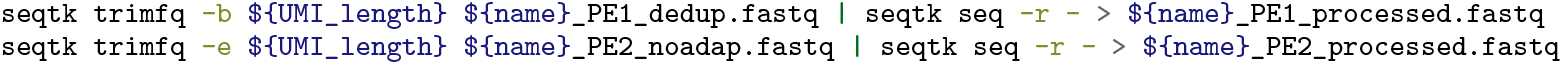

### 4.6 Remove reads aligning to rDNA

While between 70 - 80% of stable RNA is rRNA,generally less than 10% of the nascent RNA arises from rRNA. By first aligning to the rDNA, we can later estimate nascent RNA purity. Any reads that map non-uniquely to both rDNA and non-rDNA regions in the genome result in artifactual spikes at regions in the genome that share homology with the rDNA locus. Before aligning to the genome, we first align reads to rDNA and use `samtools fastq` and the `−f 0×4` flag to specify that only unmapped reads are included in the FASTQ output. We recommend the following site to help understand the meaning of samtools flags: https://broadinstitute.github.io/picard/explain-flags.html.

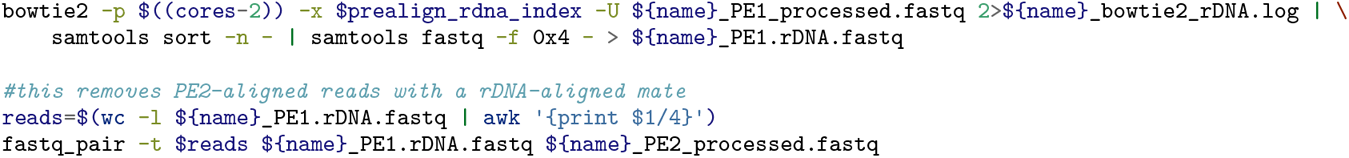

### 4.7 Genome alignment

The last processing step for individual libraries is to align to the genome. We invoke the `––rf` flag to account for the fact that we reverse complemented both reads. The samtools commands convert the file to a compressed binary BAM format and sort the reads.

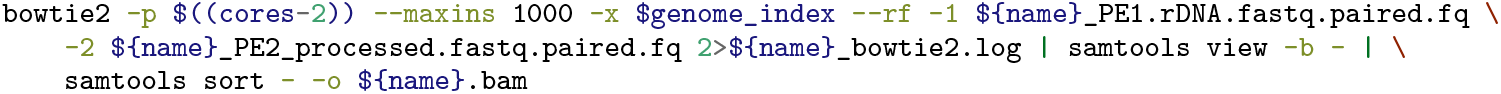

### 4.8 rDNA alignment rate

In order to calculate the rDNA alignment rate, we first count the total number of rDNA-aligned reads. Next, we use `samtools view −c −f 0×42` to count the PE1 reads that concordantly align to hg38 and not to rDNA. Lastly, we calculate the fraction of aligned reads that map to the rDNA locus and print it to the QC metrics file.

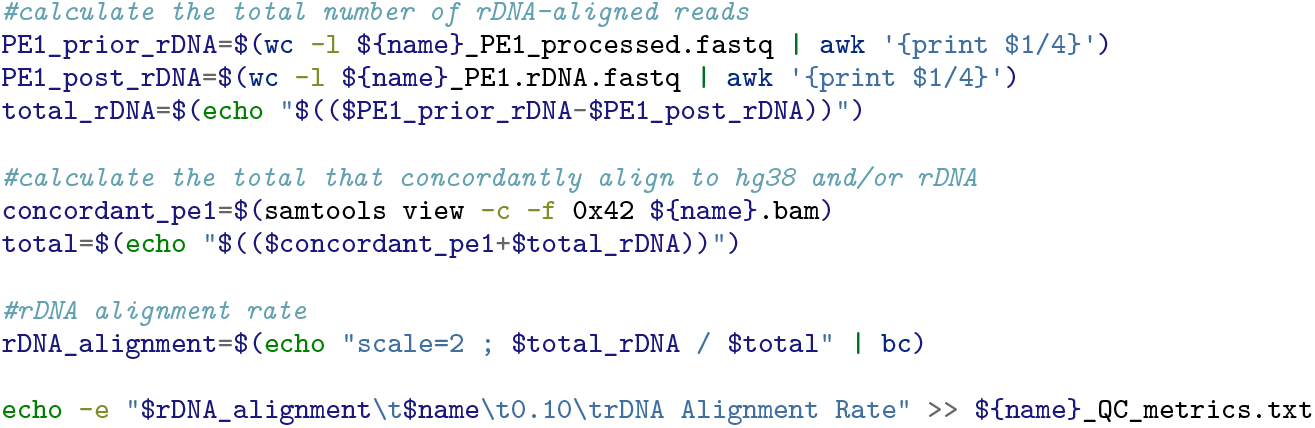

### 4.9 Mappability rate

The majority of reads should map concordantly to the genome. We expect an alignment rate above 80% for high quality libraries. As described above, we use `samtools and wc −l` to count concordantly aligned reads in the BAM alignment file and the pre-alignment FASTQ files and then divide these values to calculate the alignment rate. We found that low alignment rates typically arise from either poor quality sequencing or microorganism contamination of reagents/buffers. We recommend using FastQC to determine if the poor alignment is due to a problem with the FASTQ base quality scores ***(33)***. If the user suspects that the poor alignment is due to xenogeneic DNA contamination, we recommend using `BLAST` to query unaligned sequences to genome databases ***(34)*** The user can leverage the sequences to design PCR primers for the contaminating species and test reagents to identify the source.

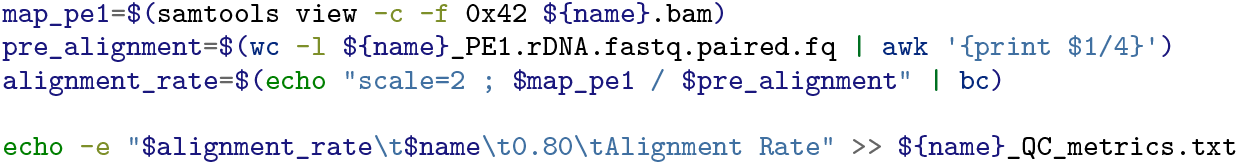

### 4.10 Complexity and theoretical read depth

The proportion of PCR duplicates in a library affects how many additional raw sequencing reads are required to achieve a target number of concordantly aligned reads. We developed `fqComplexity` to serve two purposes: 1) calculate the number of reads that are non-PCR duplicates as a metric for complexity, and 2) provide a formula and constants to calculate the theoretical read depth that will result in a user-defined number of concordant aligned reads. The proportion of reads that are PCR duplicates is related to read depth. At very low read depth nearly all reads are unique; at very high read depth the observed fraction of duplicates approaches the true PCR duplicate rate of the library. We calculate the PCR duplicate rate using the processed FASTQ file without adapter/adapter ligation products or small inserts. The FASTQ file is randomly subsampled to read depths of 10%, 20%, 30%,…,100%, and the intermediate files are deduplicated. We print the total and deduplicated counts for each subsample to the `${name}_complexity.log file`. The R script fits an asymptotic regression model and plots the model and data (Figure 2A). We recommend that at least 75% of reads are unique at a read depth of 10 million.

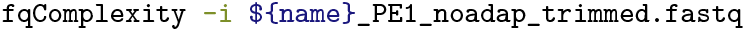

**Figure 2:**
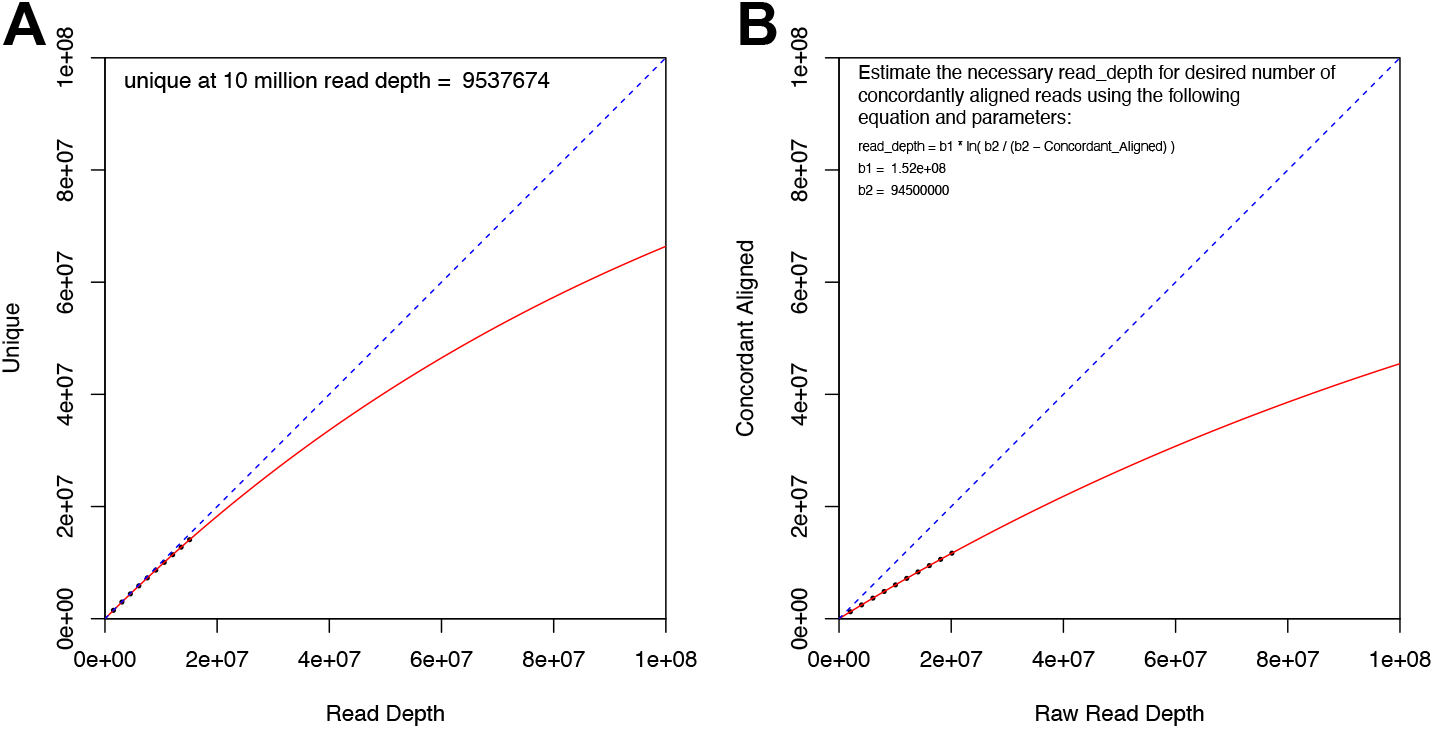
Library complexity captures information about PCR over-amplification and read depth requirements for a sample. A. We subsample the pre-processed FASTQ file to the indicated read depths (x-axis) and plot this value against the number of unique subsampled reads (y-axis). The plot includes an asymptotic regression model curve and prints the estimated number of unique reads at a read depth of 10 million. B. We use the fraction of raw PE1 reads that do not contain adapter ligation products or small inserts, the fraction of deduplicated reads that align concordantly to the non-rDNA genome, and the data from panel A to derive the theoretical read depth equation and parameters.

Sequencing depth requirements vary depending upon downstream applications and the size/gene density of the genome. We recommend three replicates and over 10 million concordantly aligned reads per replicate for differential expression analysis with human cells. Data driven approaches to define gene annotations or identify regulatory elements require higher sequencing depth. We need two factors to calculate the raw read depth necessary to achieve a specified target condordantly aligned depth. The first value is the fraction of raw PE1 reads that do not contain adapter/adapter ligation products or small inserts: `$factorX`.

The second value is the fraction of deduplicated reads that align concordantly to the non-rDNA genome: `$factorY`. Finally, we run `fqComplexity` and specify the `−x` and `−y` options to fit an asymptotic regression model to the factor-scaled log file. `fqComplexity` searches for and reuses the previous log file to avoid unnecessarily repeating subsampling and deduplication. We use the equation and constants printed in the PDF output (Figure 2B) to determine the practicality of increasing depth using the same libraries.

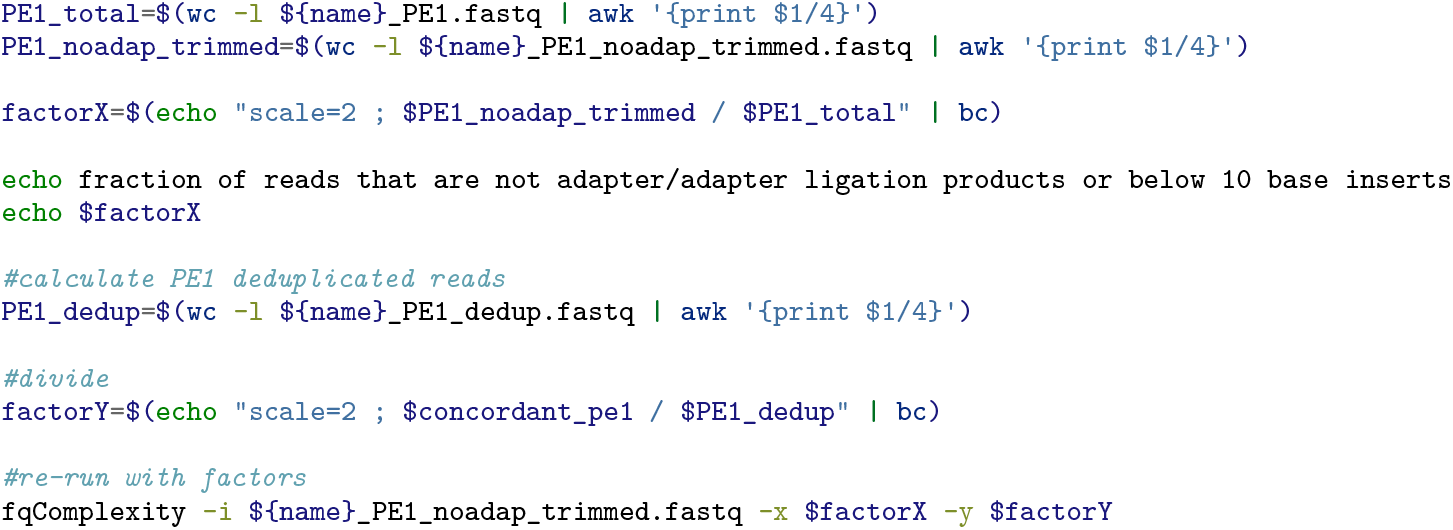

### 4.11 Run-on efficiency

RNA polymerases that are associated with gene bodies efficiently incorporate nucleotides during the run-on reaction under most conditions, but promoter proximal paused RNA polymerase require high salt or detergent to run-on efficiently ***(7, 35)***. Therefore, the pause index, or the density of signal in the promoter-proximal pause region divided by density in the gene body, is an indirect measure of run-on efficiency. Since pause windows are user-defined and variable, pause indices can differ substantially based on how they are calculated.

To determine the coverage of PRO-seq signal in genomic intervals it is convienent to convert the genomic signal to a BED6 file format. Although we are not correcting enzymatic sequence bias in this workflow, we use `seqOutBias` with the `−−no−scale` option to convert the BAM file. We include the `−−tail−edge` option to realign the end of the read so that the exact position of RNA Polymerase is specified in the BED6 output file. The `−−out−split−pairends` option separates all the paired end reads and ––stranded prints strand information in column 6.

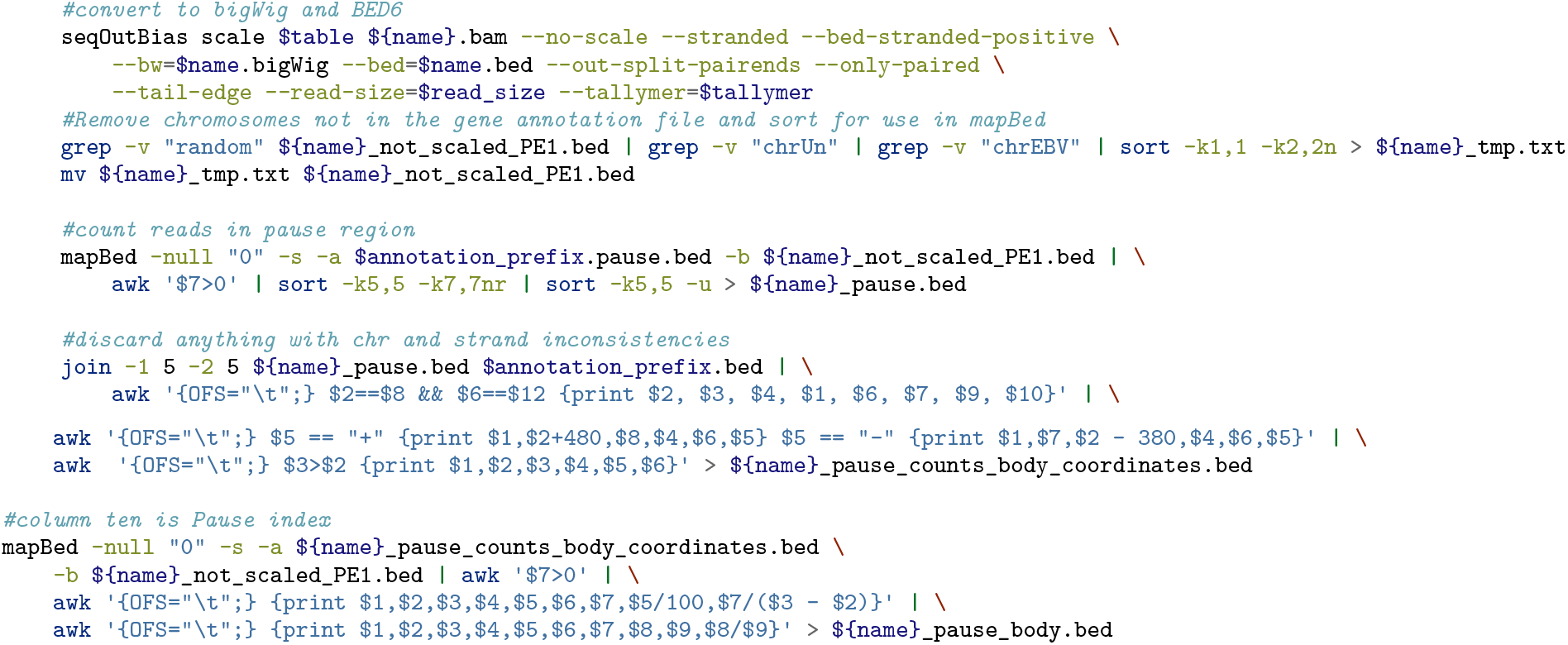

We use an R script to calculate pause indices and plot the distribution of log10 pause index values as a PDF (Figure 3).

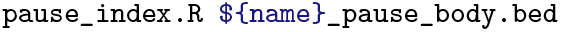

**Figure 3:**
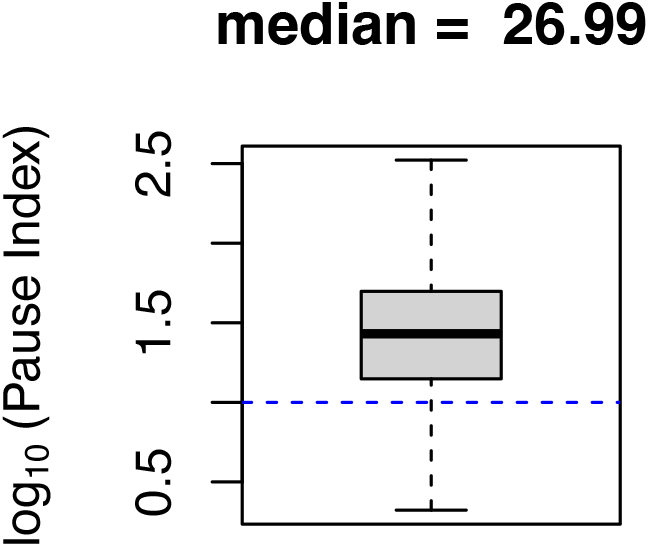
Pause index is a measure of nuclear run-on efficiency. The plot illustrates the distribution of log_10_ pause indices and includes a threshold line at a raw pause index of 10. A median pause index below 10 indicates that the library may be of poor quality.

### 4.12 Estimate nascent RNA purity with exon / intron density ratio

Exon and intron densities within each gene are comparable in nascent RNA-seq data. In contrast, RNA-seq primarily measures mature transcripts and exon density far exceeds intron density. We can infer mature RNA contamination in PRO-seq libraries if we detect a high exon density to intron density ratio. We exclude contributions from the first exon because pausing occurs in this region and artifically inflates the exon density. The distribution of log10 exon density to intron density ratios is output as a PDF (Figure 4). This metric complements rDNA alignment rate to determine nascent RNA purity.

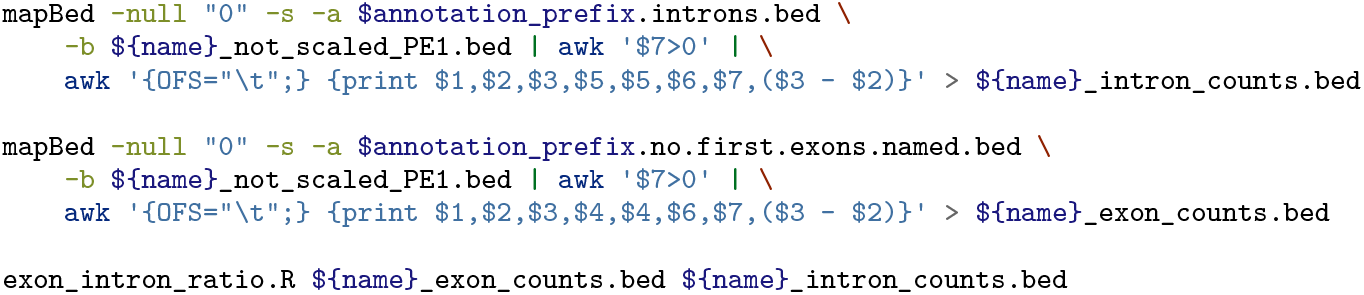

**Figure 4:**
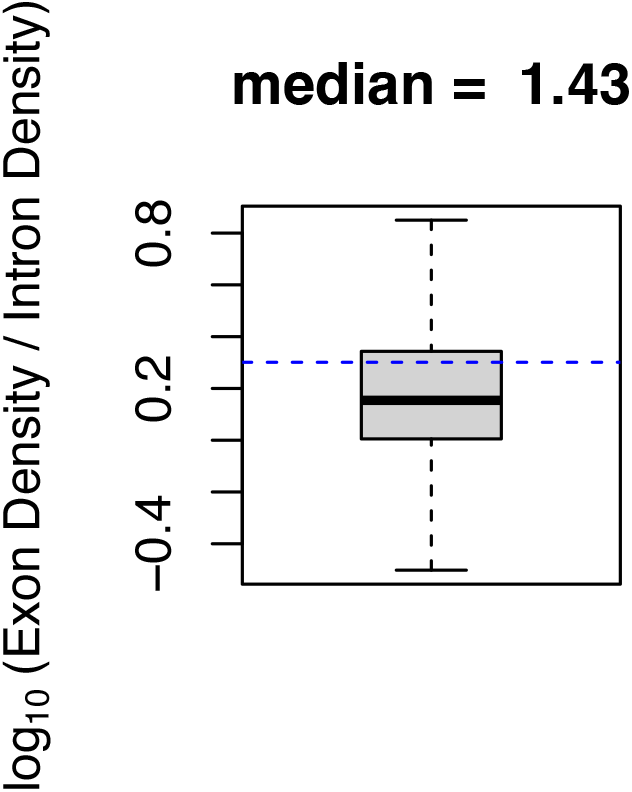
Exon density to intron density ratio is a measure of nascent RNA purity. The plot illustrates the distribution of log_10_ exon density to intron density ratios and includes a threshold line at a raw ratio of 2. A median ratio below 2 indicates mature RNA contamination is low.

### 4.13 Remove intermediate files and zip raw sequencing files

Calculating these quality control metrics necessitates many intermediate files. Many files are unused output from various processing steps or only used briefly. FASTQ files are large and rarely used in downstream analyses, so the following code chunk removes intermediate FASTQ files and compresses the original files.

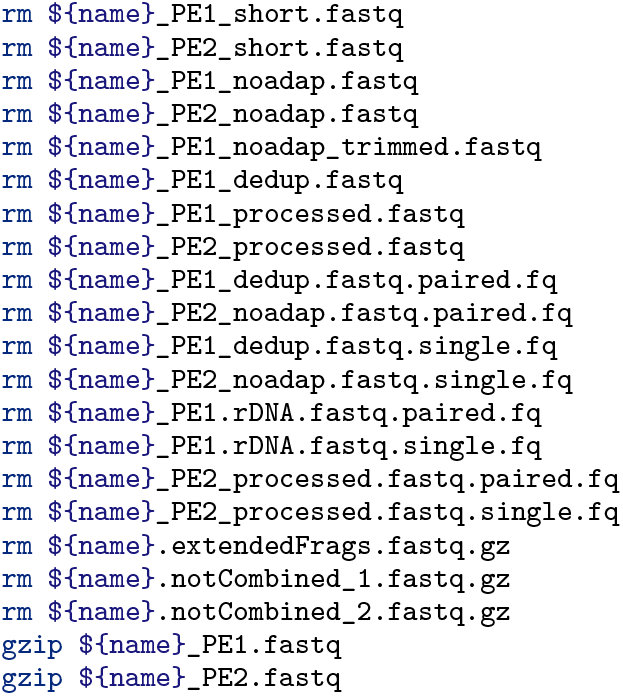

## 5 Pipeline Automation

We present the deconstructed workflow above because it is helpful to run through the code chunks individually to gain further understanding of each step. A more complete understanding of the processes allows the user to modify steps based on PRO-seq protocol variations. However, automation of routine processing and analysis is more practical once a workflow is established. Below, we provide a shell script loop that will process each set of paired end files in series. This loop can be adapted to perform all processing in parallel using a job scheduler and submission of a batch script for each set of paired end input files.

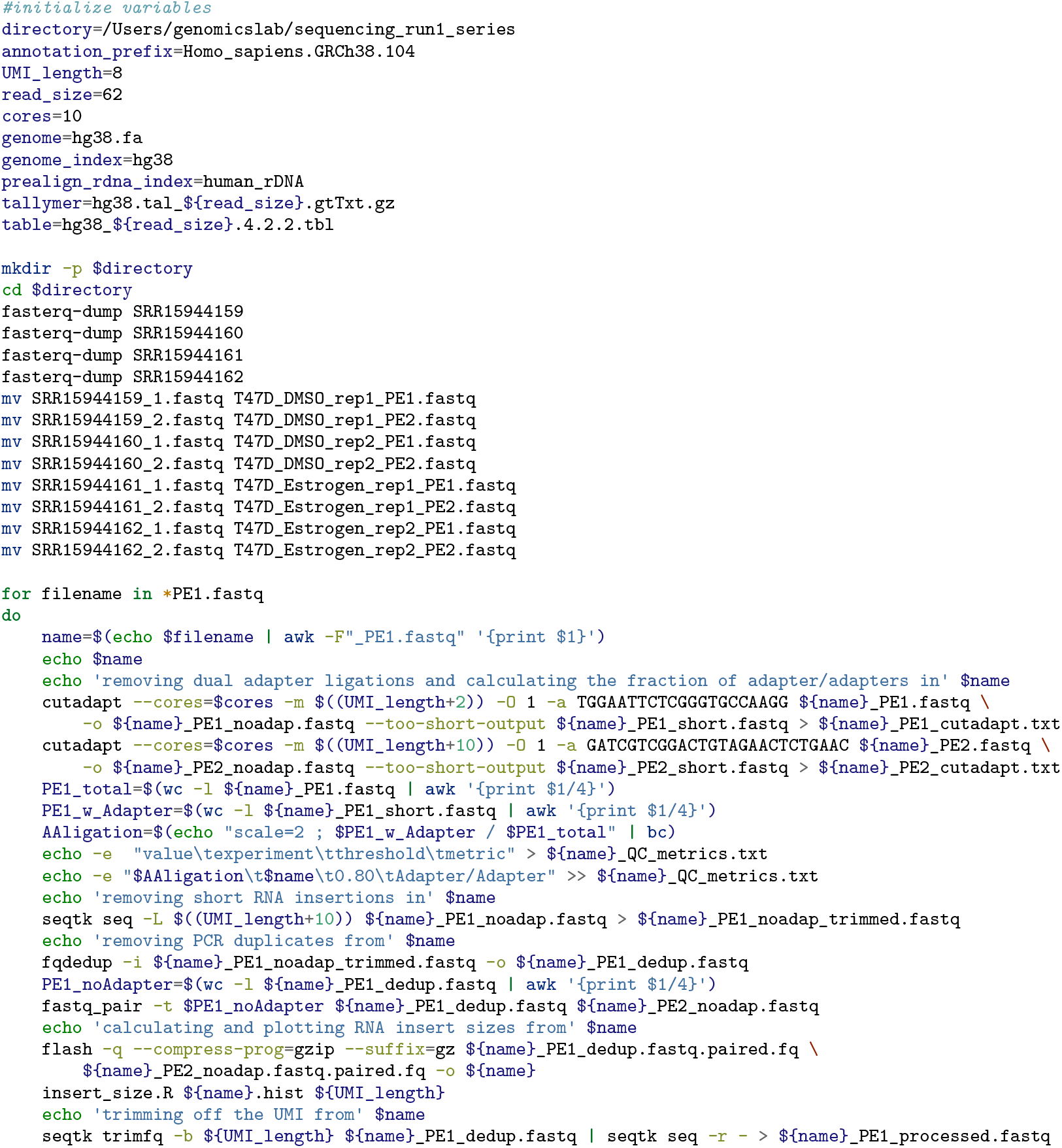
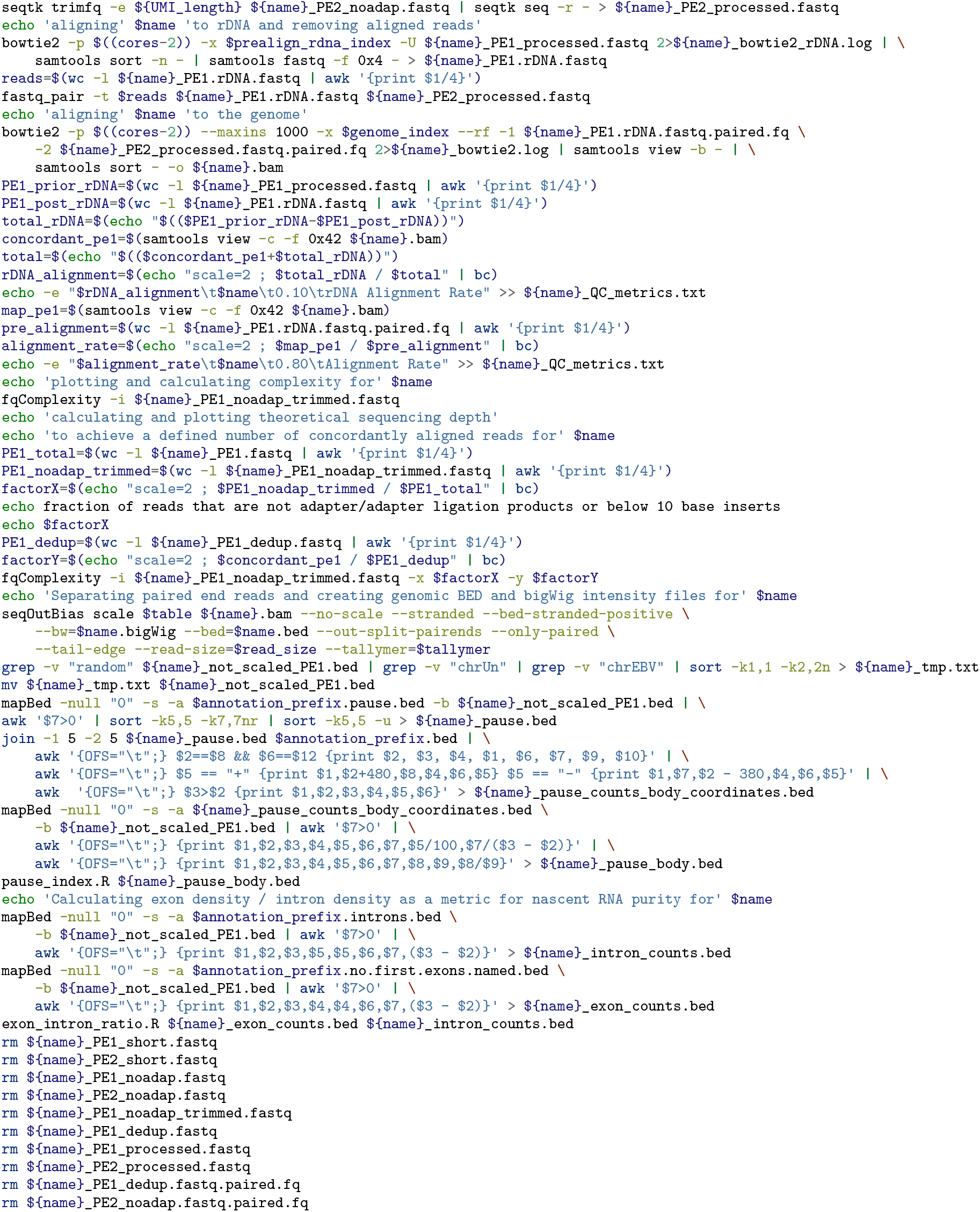
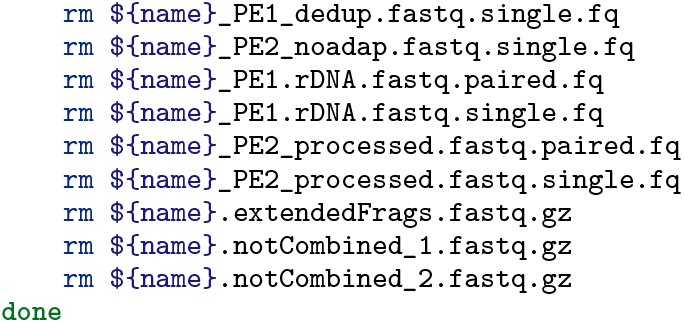

## 6 Plot all QC metrics

Individual plots for each quality control metric provide valuable information about the data, but each plot can be summarized as a single informative value. We empirically determined thresholds for each value that constitute acceptable libraries. These thresholds are not absolute and should only be used as guidelines. Below, we concatenate all the summarized metrics for the experiments and plot the results (Figure 5) and thresholds. The user can quickly glance at the plot to determine whether the quality control values fall within the acceptable range, which is shaded light green. If values are within the dark pink region, then we recommend looking back at the more detailed quality control plots to diagnose possible issues with the libraries. The user can change the term “Estrogen_treatment_PRO” to a description of their own experiment to name the output file.

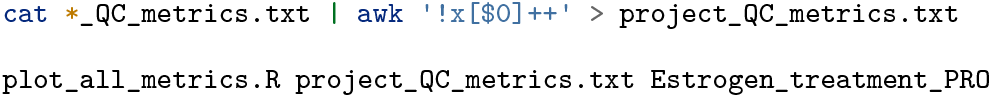

**Figure 5:**
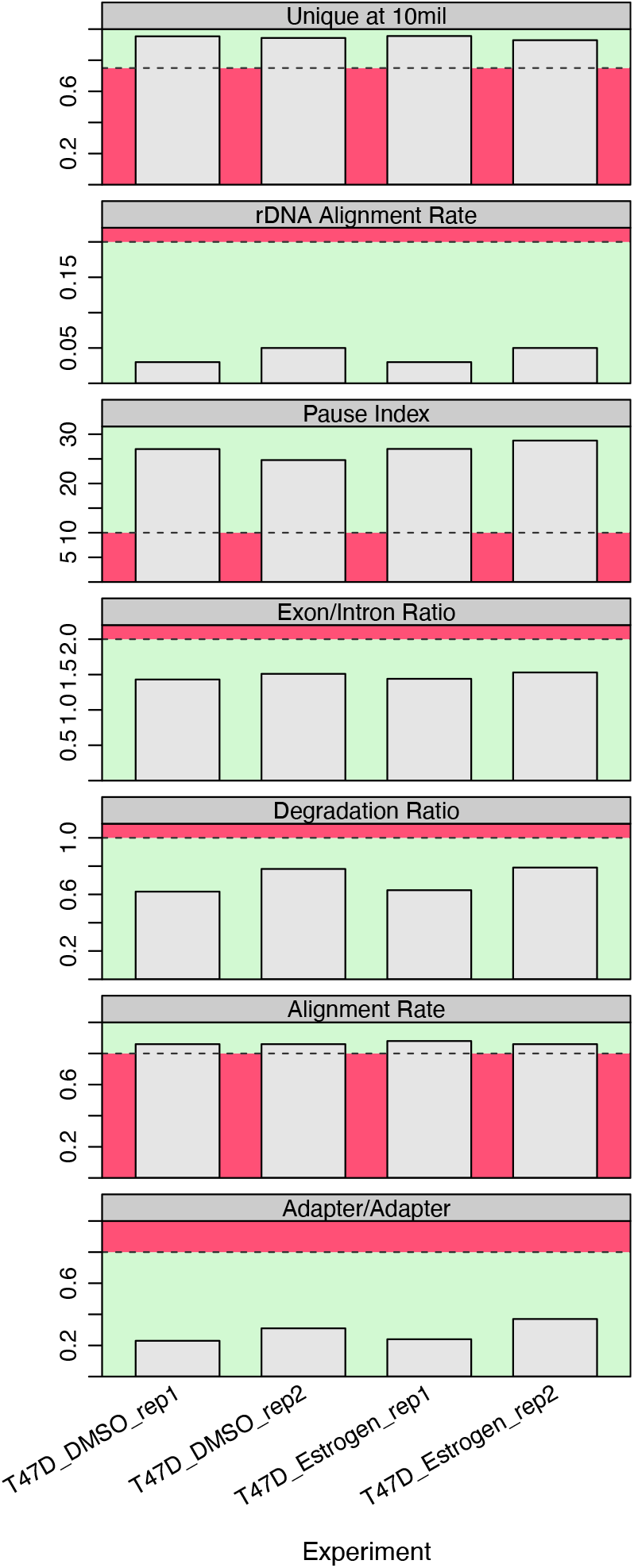
A summary plot illustrates all quality control metrics and their respective recommended thresholds. If all quality control values fall within the shaded light green range, then the libraries are likely of high quality. If values are within the dark pink region, then we recommend looking back at the more detailed quality control plots in Figures 1–4 to diagnose possible issues with the libraries.

## 7 Differential expression with DESeq2

Differential expression analysis is a common first step after routine RNA-seq and PRO-seq data processing. Below we present the `bedtools` command to count reads within gene annotations and we provide an `R` script for differential expression analysis with `DESeq2`. The script also plots the fold change between conditions and mean expression level for each gene. For simplicity we use the most upstream transcription start site and most downstream transcription termination site for annotations, but there are more accurate methods to define primary transcripts ***(5, 6)***. The `R` script requires three ordered arguments: 1) a file with the signal counts for each gene in every even row, 2) the prefix for the baseline experimental condition for which to compare (often termed “untreated”), 3) prefix name for the output PDF plot (Figure 6).

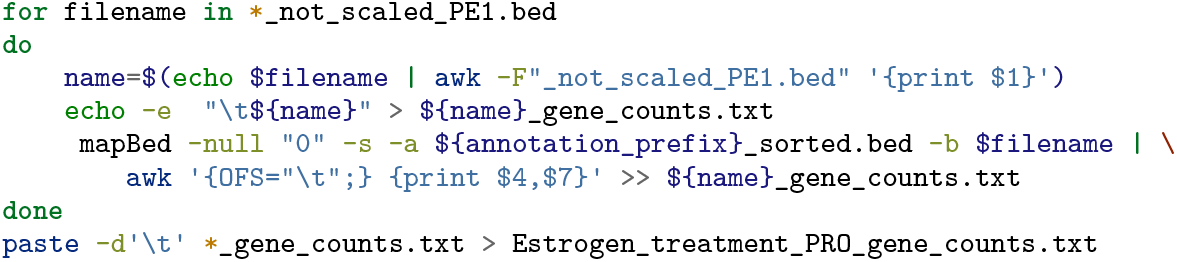

**Figure 6:**
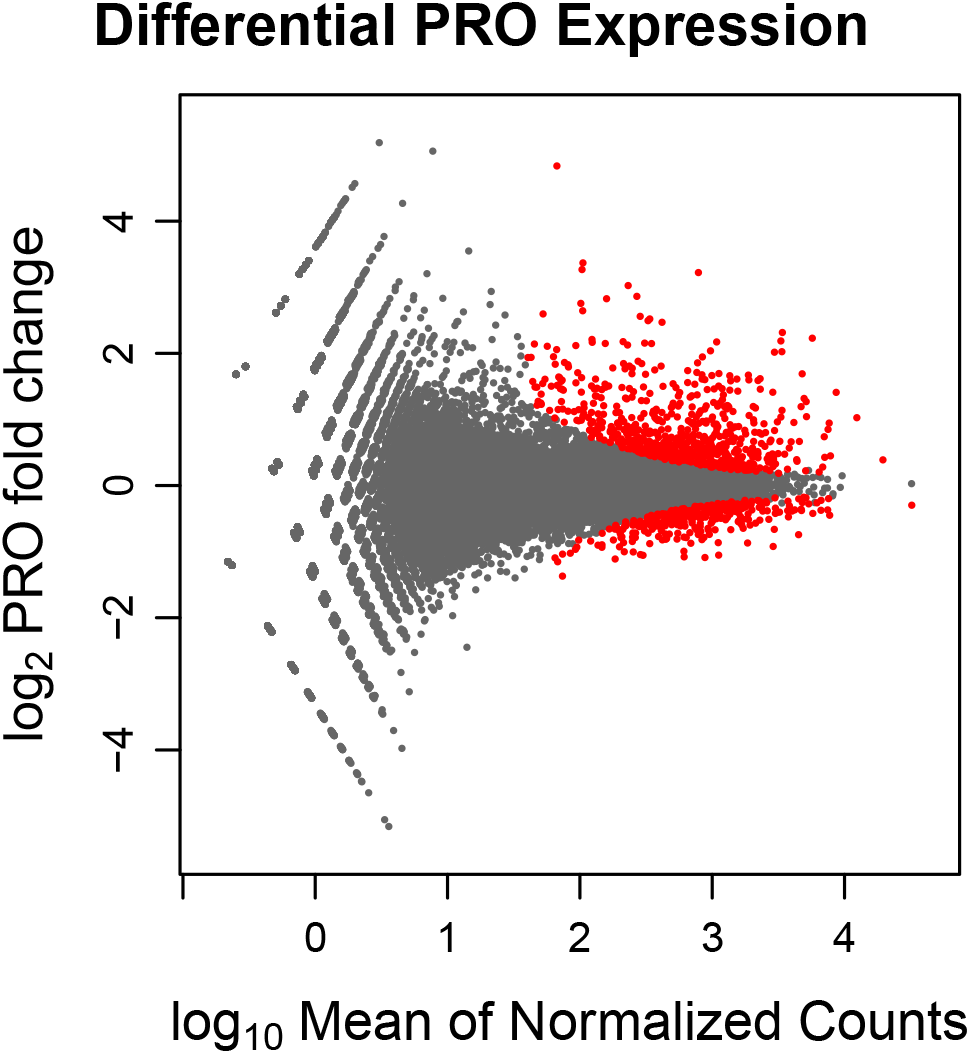
Differential expression analysis quantifies transcriptomic changes upon treating T47D cells with estrogen for an hour. Genes in red are classified as activated and repressed based on a false discovery rate of 0.05.

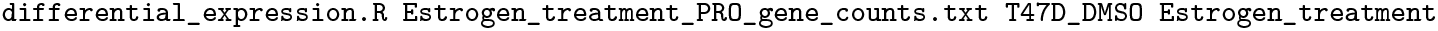

## 8 Conclusions

We provide standardized metrics and detailed plots that indicate whether libraries are of sufficiently high quality to warrant downstream analysis. The presented analyses provide information about RNA degradation, nascent RNA purity, alignment rate, library complexity, and nuclear run-on efficiency. We deconstruct each analysis and explain the biological rationale of each metric. All code and scripts are presented so that researchers can use this framework to develop their own workflows and pipelines, or as Captain Barbossa succinctly stated: “The code is more of what you’d call *guidelines* than actual rules.”

